# *Ezh2* knockout in B cells impairs plasmablast differentiation and ameliorates lupus-like disease in MRL/*lpr* mice

**DOI:** 10.1101/2022.07.21.500990

**Authors:** Xiaoqing Zheng, Mikhail G Dozmorov, Colleen E Strohlein, Sheldon Bastacky, Amr H Sawalha

**Author notes:** Please address correspondence to Amr H. Sawalha, MD. Address: 7123 Rangos Research Center, 4401 Penn Avenue, Pittsburgh, PA 15224, USA.

## Abstract

**Objectives:** Enhancer of zeste homolog 2 (EZH2) regulates B cell development and differentiation. We have previously demonstrated increased EZH2 expression in peripheral blood mononuclear cells isolated from lupus patients. The goal of this study was to evaluate the role of B cell EZH2 expression in lupus pathogenesis.

**Methods:** We generated an MRL/*lpr* mouse with floxed *Ezh2*, which was crossed with CD19-Cre mice to examine the effect of B cell EZH2 deficiency in MRL/*lpr* lupus-prone mice. Differentiation of B cells was assessed by flow cytometry. Single cell RNA sequencing and single cell B cell receptor sequencing were used to investigate compositional and functional changes of B cell subsets. *In vitro* B cell culture with an XBP1 inhibitor was performed. EZH2 and XBP1 mRNA levels in CD19^+^ B cells isolated from SLE patients and healthy controls were analyzed.

**Results:** We show that *Ezh2* deletion in B cells significantly decreased autoantibody production and improved glomerulonephritis. B cell development was altered in the bone marrow and spleen of EZH2-deficient mice. Differentiation of plasmablasts was impaired. Single cell RNA sequencing showed that XBP1, a key transcription factor in B cell development, is downregulated in the absence of EZH2. Inhibiting XBP1 *in vitro* impairs plasmablast development similar to EZH2-deficient mice. Single cell B cell receptor RNA sequencing revealed defective immunoglobulin class switch recombination in EZH2-deficient mice. In human lupus B cells, we observed a strong correlation between EZH2 and XBP1 mRNA expression levels.

**Conclusion:** EZH2 overexpression in B cells contributes to disease pathogenesis in lupus.

## Introduction

Systemic lupus erythematosus (SLE or lupus) is a multi-system autoimmune disease characterized by B cell autoreactivity and autoantibody production. Substantial research has pointed to the involvement of both genetic and epigenetic mechanisms in the pathogenesis of lupus [1, 2]. In particular, abnormal DNA methylation has been shown to play crucial roles in this disease [3-5]. We have previously demonstrated that the expression of the key epigenetic modulator Enhancer of zeste homolog 2 (EZH2) is dysregulated in lupus peripheral blood mononuclear cells compared to healthy controls [6]. EZH2, a core component of polycomb repressive complex 2 (PRC2), is a histone methyltransferase that catalyzes di- and tri-methylation of lysine 27 in histone H3 (H3K27) [7]. We have shown that overexpression of EZH2 in lupus CD4+ T cells is associated with pro-inflammatory epigenetic changes with potential pathogenic consequences in lupus patients [5, 8, 9]. Further, we demonstrated that inhibiting EZH2 ameliorates lupus-like disease in animal models [6], which was subsequently confirmed by others [10, 11]. However, the role of EZH2 and EZH2-regulated genes in lupus B cells has not been adequately explored.

EZH2 plays a critical role in B cell development and differentiation [12]. EZH2 is required for germinal center (GC) B cell development and function, and EZH2 gain of function leads to GC hyperplasia [13]. Spontaneous and expanded GCs are a feature of murine and human lupus [14]. EZH2 regulates immunoglobulin heavy chain rearrangement in progenitor B (Pro-B) cells [15], and represses immunoglobulin κ-chain complex recombination in precursors of B (Pre-B) cells [16]. Further, EZH2-dependent regulation of transcriptional activity and cellular metabolism are required for antibody production [17].

We have previously shown that EZH2 is overexpressed in lupus B cells compared to normal healthy controls [6]. This observation was independently confirmed [18], with additional data demonstrating correlation between EZH2 expression levels in B cells with disease activity and autoantibody production in lupus patients [18]. Activation of Syk and mTORC1 synergistically induce EZH2 expression in B cells [18]. Indeed, both Syk and mTORC1 are activated in B cells from lupus patients, providing a mechanism explaining EZH2 overexpression in lupus B cells [18, 19].

In this study, we generated CD19-Cre *Ezh2*^*fl/fl*^ mice on lupus-prone MRL/*lpr* background to investigate the pathogenic role of EZH2 in lupus B cells. Our results showed that *Ezh2* deletion in B cells significantly decreased autoantibody production and alleviated lupus glomerulonephritis. Differentiation of antibody secreting cells and class switch recombination were significantly impaired in EZH2-deficient mice. B-cell specific EZH2 deficiency led to reduced expression of XBP1, which is a key transcription factor involved in plasmablast/plasma cell development and immunoglobulin production. Further, in human lupus B cells, the mRNA expression of EZH2 was very strongly correlated with XBP1 expression.

## Methods

*Mice*. An *Ezh2*^*fl/fl*^ mouse on the MRL/*lpr* lupus-prone background was generated by genetic engineering using CRISPR/Cas9 technology at the Innovative Technologies Development Core and the Mouse Embryo Services Core (Department of Immunology, University of Pittsburgh School of Medicine). LoxP/LoxP alleles were flanked on *Ezh2* exon 17, whose deletion mediated by Cre will cause a frameshift and ultimately an absence of transcription (**Figure 1A**). CD19-Cre MRL/*lpr* mice were a gift from Dr. Mark Shlomchik at the University of Pittsburgh [20]. CD19-Cre MRL/*lpr* mice were crossed with *Ezh2*^*fl/fl*^ mice, then heterozygous CD19-Cre *Ezh2*^*fl/wt*^ mice were bred with *Ezh2*^*fl/fl*^ mice. CD19-Cre *Ezh2*^*fl/fl*^ mice were used as the experimental mouse group and littermate *Ezh2*^*fl/fl*^ lacking the Cre transgene as controls. Genotyping primers were as follows: *Ezh2* F1: GAACAAGGGGTTCGGGATCA; *Ezh2* R1: AGCACACCCACTTACACAGG. *CD19* forward primer: AATGTTGTGCTGCCATGCCTC; *CD19-Cre* reverse primer: TTCTTGCGAACCTCATCAC; *CD19* reverse primer: GTCTGAAGCATTCCACCGGAA. Q5 high-fidelity DNA polymerases (NEW ENGLAND BioLabs) were used for genotyping (**Supplementary Methods**). DNA (DNeasy Blood & Tissue Kit, QIAGEN) isolated from CD19^+^ B cells (stained with brilliant violet (BV) 421 anti-mouse CD19 antibody, Clone 6D5, BioLegend) sorted by FACS Arial II (BD Bioscience) were used to measure deletion efficiency of *Ezh2*. The primers used were *Ezh2* F2: GTGTTCTAACAGTCTTGACCA; *Ezh2* R1: AGCACACCCACTTACACAGG (as shown in **Figure 1A**). Beta-actin primers used were *Actb* F1: CATTGCTGACAGGATGCAGAAGG; *Actb* R1: TGCTGGAAGGTGGACAGTGAGG. CD19-Cre *Ezh2*^*fl/fl*^ experimental mice and *Ezh2*^*fl/fl*^ control mice were housed under the same pathogen-free conditions for 24 weeks before they were sacrificed. *Ezh2* deletion was confirmed with DNA isolated from CD19^+^ splenocytes in the CD19-Cre *Ezh2*^*fl/fl*^ compared to the *Ezh2*^*fl/fl*^ littermate control mice (**Figure 1B**).

**Figure 1.**
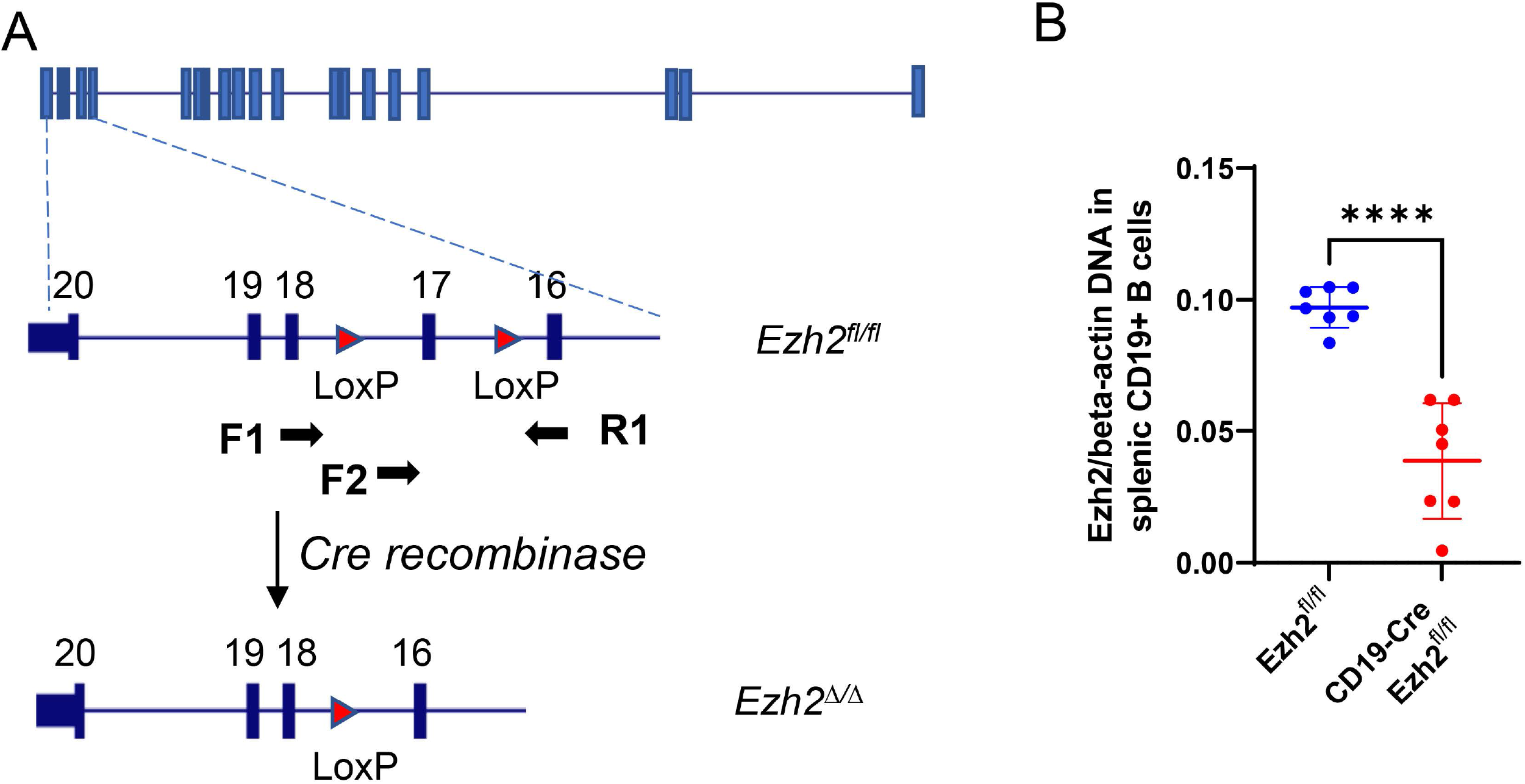
CD19^+^ *Ezh2* knockout model in MRL/*lpr* mice. (**A**) *Ezh2* genomic targeting strategy in MRL/*lpr* mice. Exon 17 in the SET methyltransferase domain of *Ezh2* was flanked by LoxP sites to be deleted by Cre recombinase. F1, F2, and R1 indicate primers loci. F1-R1 primers were used for *Ezh2*^*fl/fl*^ genotyping and F2-R1 primers were to check *Ezh2* deletion efficiency. (**B**) DNA *Ezh2* measured with F2-R1 primers in qPCR normalized to *beta-actin* in CD19^+^ B cells sorted with flow cytometry. Data are presented as mean ± SD, n=7, \*\*\*\**p*<0.0001, unpaired two-tailed *t* test.

### Flow cytometry analysis of bone marrow and spleen B cells

Bone marrow (BM) cells were isolated from both the femur and tibia. The spleen was mashed and prepared into single-cell suspension. Red bloods cells (RBC) from BM and spleen were lysed with RBC lysis buffer (Invitrogen) in room temperature for 5 minutes. Cells were resuspended in PBS containing 1% FBS and incubated with mouse FcR blocking reagent (Miltenyi Biotec) for 10 minutes at 4°C before antibody staining with 1:50 dilution for 30 minutes at 4°C. Flow cytometry data were acquired by LSRFortessa (BD Bioscience) and analyzed by FlowJo software (version 10.8, BD Bioscience).

The following antibodies were used to detect B cell populations in the BM: brilliant violet (BV) 421 anti-mouse CD19 antibody (Clone 6D5, BioLegend), PE/Cyanine5 anti-mouse/human CD45R/B220 (Clone RA3-6B2, BioLegend), PE-Dazzle 594 rat anti-mouse CD43 (Clone 1B11, BioLegend), VioBright 515 anti-mouse IgM (Clone REA979, Miltenyi Biotec), SuperBright 702 rat anti-mouse IgD (Clone 11-26c, eBioscience), and fixable near-IR dead cells stain kit (Invitrogen).

To analyze B cell subsets in the spleen, the following antibodies were used: BV786 rat anti-mouse CD93 (Clone 493, BD Bioscience), BV480 rat anti-mouse CD21/CD35 (Clone 7G6, BD Bioscience), BV605 anti-mouse CD23 (Clone B3B4, BD Bioscience), PerCP-eFluor 710 anit-GL7 (Clone GL7, eBioscience), APC anti-mouse CD38 (Clone 90, BioLegend), PE/Cyanine7 anti-mouse CD138 (Syndecan-1) (Clone 281-2, BioLegend), and the same anti-CD19, anti-B220, anti-IgM, anti-IgD, and live/dead staining used for BM cells.

### Plasma autoantibody measurement

Plasma samples were collected from mice at 24 weeks of age. Anti-double-stranded DNA (anti-dsDNA) antibody was measured by ELISA following the manufacturer’s instructions (Alpha Diagnostic International, San Antonio, USA).

### Assessment of proteinuria and nephritis

Urine was collected within one week before mice were sacrificed. Urine albumin and creatinine were measured using ELISA kits (Alpha Diagnostic International, San Antonio, USA, and R&D systems, Minneapolis, USA, respectively). Urine samples were diluted to fit within the standards range of each kit. Urine albumin-to-creatinine ratio (UACR) was calculated to assess proteinuria. Kidneys were harvested, fixed with formalin for 2 days, then dehydrated in 70% ethanol and embedded in paraffin. Sections were cut and stained with hematoxylin and eosin (H&E). Glomerulonephritis and interstitial inflammation were reviewed by a renal pathologist in a blinded manner. Glomerulonephritis was scored on a scale from 1 to 6, and interstitial inflammation was scored on a scale from 0 to 4, as previously described [21].

### Single cell RNA sequencing (scRNA-seq)

Splenocytes isolated from four female mice (two *Ezh2*^*fl/fl*^ and two CD19-Cre *Ezh2*^*fl/fl*^) were stained with BV421 anti-mouse CD19 antibody (Clone 6D5, BioLegend) and live/dead dye, and sorted with FACS Arial II (BD Bioscience). Splenic B cells from each mouse were incubated with a unique TotalSeq-C0301 anti-mouse hashtag antibody conjugated to a Feature Barcode oligonucleotide (BioLegend) to label each mouse at 4°C in 100 μl PBS+0.04% BSA for 30 minutes. Splenic B cells from four mice were pooled and gel beads in emulsion were formed. Single cell 5’ gene expression sequencing library, single cell V(D)J enriched library for B cell receptor (BCR) sequencing, and cell surface protein Feature Barcode library were prepared following 10X Genomics instructions in the Single Cell Core, University of Pittsburgh. Sequencing was performed with 100-cycle NovaSeq v1.5 to achieve a total of 800M reads, including ∼5000 reads per cell for the BCR library and 5000 reads for Feature Barcode library and the rest for gene expression library.

### Single-cell RNA sequencing analysis

Cell Ranger v. 6.0.1 (10X Genomics, Pleasanton, CA, USA) was used to process raw sequencing data. Data analysis was performed using Seurat v.4.1 R package. Integration of two *Ezh2*^*fl/fl*^ and two CD19-Cre *Ezh2*^*fl/fl*^ datasets was performed as described [22] (“Introduction to scRNA-seq integration” vignette) with minor modifications. All genes were used for integration. 11,356 cells were analyzed from two female *Ezh2*^*fl/fl*^ mice with an average of 21,318 reads per cell and 1,652 median genes per cell. 4,971 cells were sequenced and analyzed from two female CD19-Cre *Ezh2*^*fl/fl*^ mice with an average of 10,808 reads per cell and 1,211 median genes per cell. The standard workflow for visualization and clustering was used (“Seurat - Guided Clustering Tutorial” vignette). Clustering resolution (“resolution” parameter in the FindClusters function) was set to 0.5 to obtain biologically expected number of clusters. Cluster-specific conserved markers were detected using the FindConservedMarkers function. Clusters were manually annotated based on cluster-specific differentially expressed genes; selected clusters were merged to represent major cell types (“GC cells,” “Plasmablast,” “Plasma cells”). B-cell receptor sequencing data for each condition were annotated with cluster-specific barcodes defined at the RNA-seq analysis step. “RBC, DCs and monocytes,” “T cells and NK cells,” “Neutrophil,” and “RBC” clusters were removed. Immunoglobulin heavy chain (IGH) data from filtered contig annotations were selected, and the proportions of the constant region gene family per cluster were quantified. Plots were made using Seurat’s visualization functions or the ggplot2 v3.3.5 R package. All computations were performed in R/Bioconductor v.4.1.0 [23].

### In vitro XBP1 inhibition in cultured B cells

Splenic B cells from 24-week old MRL/*lpr* mice were isolated with a Pan B Cell Isolation Kit (Miltenyi Biotec, Bergisch Gladbach, Germany). B cells were cultured with RPMI-1640 medium (Lonza, Basel, Switzerland) supplemented with 10% fetal bovine serum (FBS, Life Technologies, Carlsbad, CA, USA), 100 μg/ml penicillin-streptomycin (Sigma-Aldrich, St. Louis, Missouri, USA), 2 mM L-glutamine (Gibco, Waltham, MA USA) and 50 μM 2-Mercaptoethanol (Sigma-Aldrich, St. Louis, Missouri, USA), and with 3 μg/ml TLR7 agonist imiquimod (InvivoGen, San Diego, CA, USA) for 3 days. Imiquimod-stimulated B cells were treated simultaneously with or without the XBP1 inhibitor 4μ8c [24] (20 μM, Sigma-Aldrich, St. Louis, Missouri, USA) dissolved in DMSO (Sigma-Aldrich, St. Louis, Missouri, USA) for 3 days.

### Analysis of CD19^+^ B cells from human SLE patients and healthy controls

*EZH2* and *XBP1* mRNA levels in CD19^+^ B cells isolated from female SLE patients and female age-matched healthy controls were downloaded from GEO database (accession number GDS4185) [25, 26]. *Statistics*. Data were reported as mean ± SD or median ± interquartile range. To compare two groups, 2-tailed Student’s *t* test or Mann-Whitney *U* test was used as indicated. In single cell sequencing data, Fisher’s exact test of normalized cell frequencies after excluding non-B cell clusters was used. Statistical analysis was performed using GraphPad Prism 9.2.0 (GraphPad, San Diego, CA, USA).

### Study approval

This study was approved by the University of Pittsburgh Institutional Animal Care and Use Committee (Mouse protocol #19085735).

## Results

### B cell-specific Ezh2 deletion decreases autoantibody production in MRL/lpr mice

Plasma anti-dsDNA antibody levels were significantly lower in CD19-Cre *Ezh2*^*fl/fl*^ mice compared to *Ezh2*^*fl/fl*^ littermate controls (**Figure 2A**). Female *Ezh2*^*fl/fl*^ MRL/*lpr* mice had high anti-dsDNA antibodies level compared to male *Ezh2*^*fl/fl*^ MRL/*lpr* mice, and *Ezh2* deletion in B cells significantly decreased anti-dsDNA levels in female but not male mice (**Supplementary Figure 1A**). These results indicate that EZH2 is required to produce high levels of anti-dsDNA antibodies in MRL/*lpr* mice, and that EZH2 deficiency in B cells significantly repressed autoantibody production in this lupus-prone mouse model.

**Figure 2.**
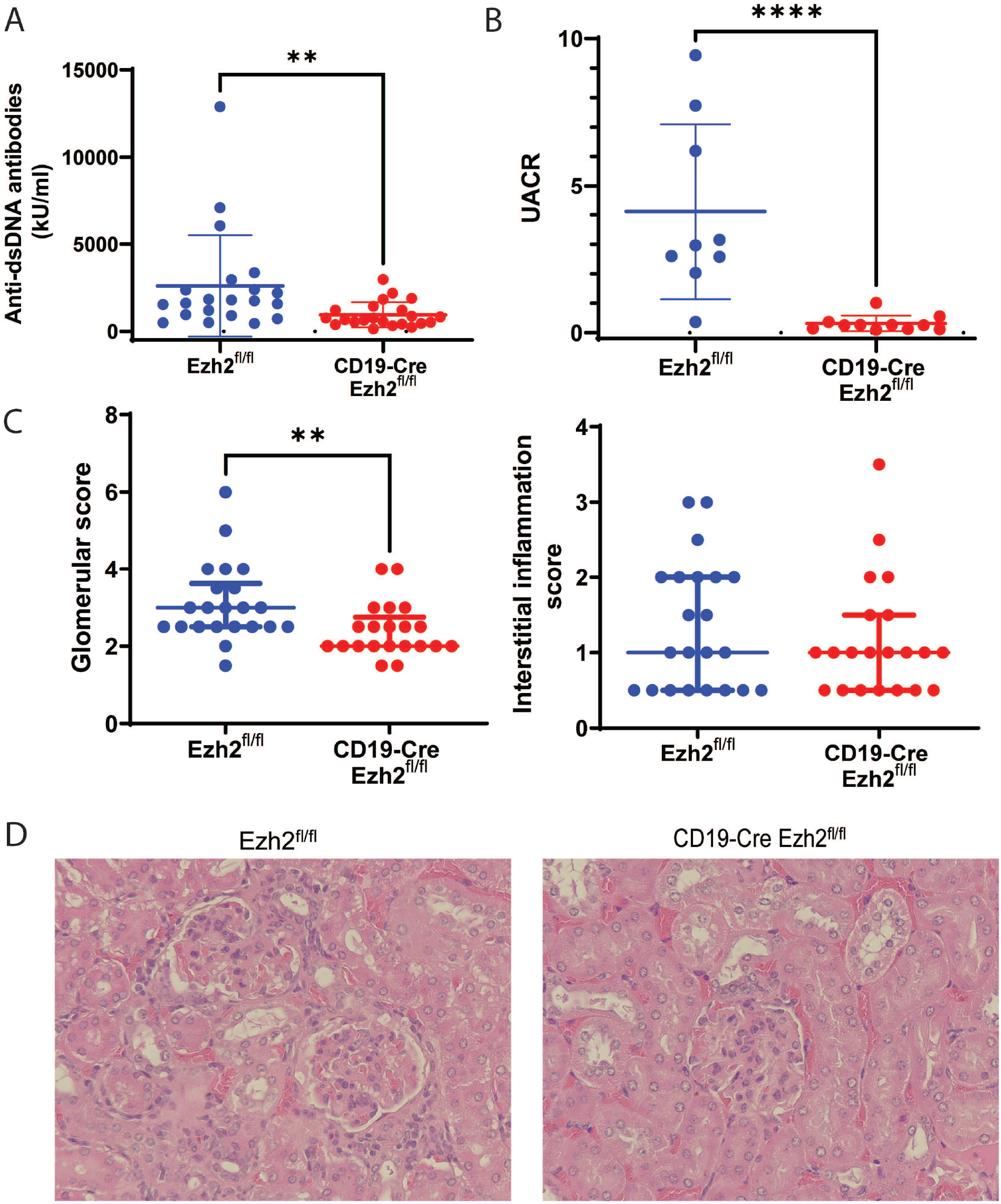
EZH2 deficiency in CD19^+^ B cells decreases serum autoantibody levels and ameliorates lupus nephritis. **A)** Serum anti-double stranded (ds) DNA antibody levels (n=21 and n=22, in *Ezh2*^*fl/fl*^ and CD19-Cre *Ezh2*^*fl/fl*^ mice, respectively), **(B)** urine albumin to creatinine ratio (UACR) (n=9 and n=11, in *Ezh2*^*fl/fl*^ and CD19-Cre *Ezh2*^*fl/fl*^ mice, respectively), **(C)** glomerular scores and interstitial inflammation scores (n=22 and n=21, in *Ezh2*^*fl/fl*^ and CD19-Cre *Ezh2*^*fl/fl*^ mice, respectively), and **(D)** representative hematoxylin and eosin (H&E) images in the female *Ezh2*^*fl/fl*^ and CD19-Cre *Ezh2*^*fl/fl*^ mice. Data in (**A**) and (**B**) are shown as mean ± SD, data in (**C**) are presented as median with interquartile range, ** *p*<0.01, \*\*\*\**p*<0.0001, two-tailed Mann-Whitney test. Magnification in (**D**), 40X.

### Ezh2 deletion in B cells alleviates lupus nephritis

Since anti-dsDNA antibody levels were significantly decreased in CD19-Cre *Ezh2*^*fl/fl*^ mice, we examined the effect of B cell-*Ezh2* deletion on lupus nephritis in MRL/*lpr* mice. Urine albumin to creatinine ratio (UACR) was used to evaluate renal disease severity. UACR was significantly lower in CD19-Cre *Ezh2*^*fl/fl*^ mice compared to littermate controls (**Figure 2B**), both in female and male mice (**Supplementary Figure 1B**). In addition, mice kidneys were fixed, sectioned, stained with H&E and blindly reviewed by a renal pathologist.

Glomerulonephritis scores were significantly lower in CD19-Cre *Ezh2*^*fl/fl*^ mice compared to *Ezh2*^*fl/fl*^ mice, but not interstitial nephritis scores (**Figure 2C, supplementary Figure 1C and 1D**). Representative kidney H&E staining images from CD19-Cre *Ezh2*^*fl/fl*^ and control mice are shown in **Figure 2D**. *Ezh2* deletion in B cells had sex-specific effects on peripheral lymphoproliferation, males had significantly larger spleens and similar size of lymph nodes, while females had a trend of smaller spleens and significantly smaller lymph nodes compared to *Ezh2*^*fl/fl*^ mice (**Supplementary Figure 2**). There was no significant difference of survival and skin lesions between CD19-Cre *Ezh2*^*fl/fl*^ and control mice (data not shown).

### Ezh2 deletion in B cells influences B cell development and differentiation in the bone marrow and spleen

B cell lymphopoiesis starts in the bone marrow and matures in secondary lymphoid organs, such as the spleen, with linear and stepwise processes as shown in **Figure 3A**. Genes for immunoglobin (Ig) heavy chain and light chain are rearranged at the Pro-B and Pre-B stages, respectively. With cell surface IgM (sIgM) expressed, immature B cells are formed, which will egress to peripheral lymph organs as transitional B cells. In the spleen, B cells will mature and develop into marginal zone (MZ) B cells with poor BCR reactivity, or follicular (FO) B cells with intermediate BCR reactivity [27]. Encountered with antigens, MZ B cells will differentiate into plasmablasts (PBs), and produce low- and high-affinity antibodies, as well as memory B cells [27]. FO B cells will become germinal center (GC) B cells and differentiate into memory B cells or PB/plasma cells (PCs) and produce high-affinity antibodies [28].

**Figure 3.**
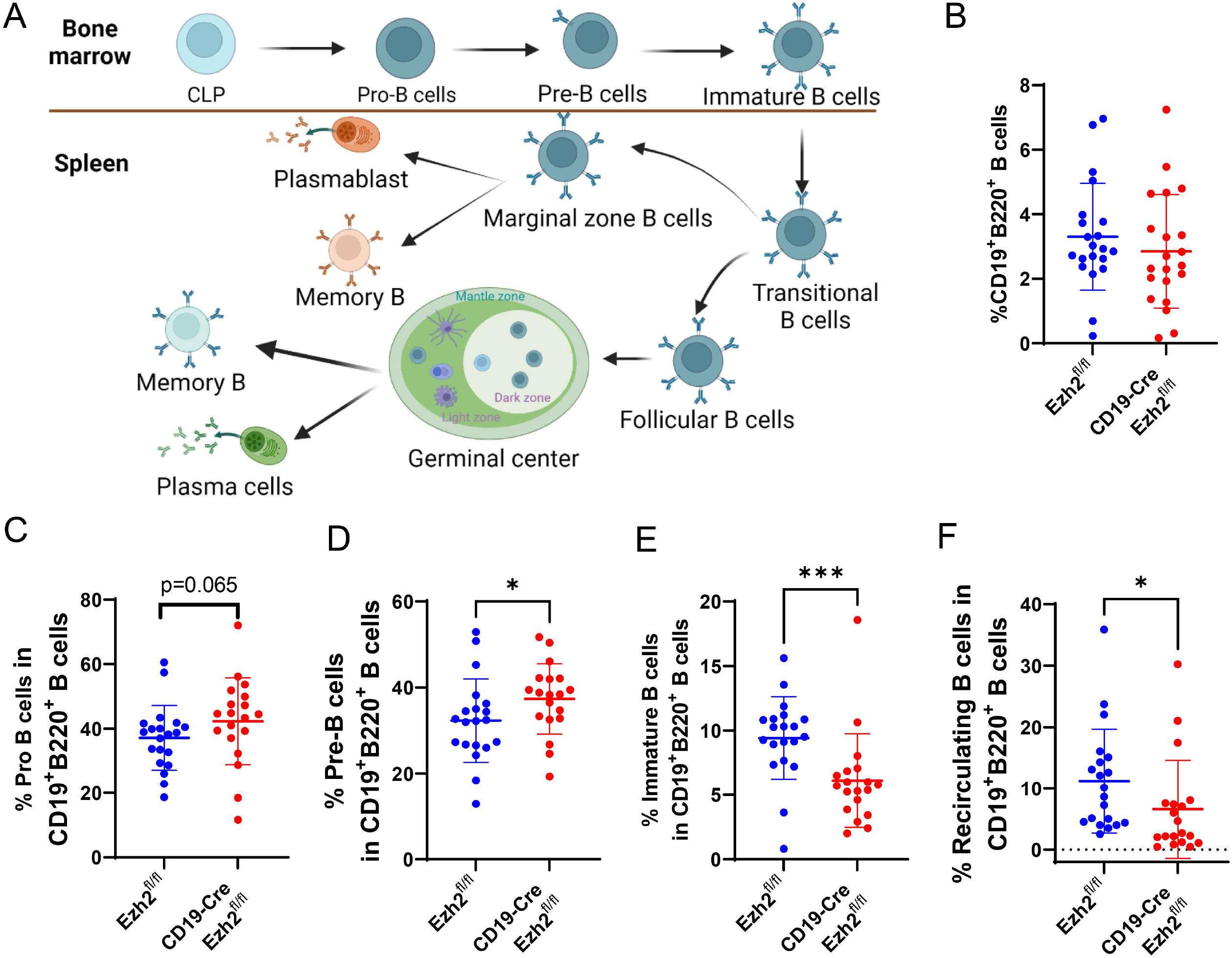
EZH2 deficiency negatively affects B cells development in the bone marrow. **(A)** A schematic representation of B cell development and differentiation in the bone marrow and spleen. The figure was created using Biorender.com. Frequency of **(B)** CD19^+^B220^+^ B cells, and proportion of **(C)** Pro-B cells, **(D)** Pre-B cells, **(E)** immature B cells, and **(F)** recirculating B cells in CD19^+^B220^+^ B cells in *Ezh2*^*fl/fl*^ (n=21/20) and CD19-Cre *Ezh2*^*fl/fl*^ (n=21/19) mice. CLP, common lymphoid progenitor. Data are shown as mean ± SD. * *p*<0.05, *** *p*<0.001, two-tailed Mann-Whitney test.

We found that the proportion of total CD19^+^B220^+^ B cells in the bone marrow (BM) was not significantly different in CD19-Cre *Ezh2*^*fl/fl*^ mice compared to *Ezh2*^*fl/fl*^ (**Figure 3B**). Gating strategies are shown in **Supplementary Figure 3**. Focusing on B cell subsets, Pro-B cells showed a trend for being increased (**Figure 3C**), Pre-B cells were significantly increased (**Figure 3D**), whereas immature B cells (**Figure 3E**) and recirculating B cells (**Figure 3F**) were significantly decreased in CD19-Cre *Ezh2*^*fl/fl*^ mice compared to littermate controls. Male and female mice exhibited similar changes in BM B cells (**Supplementary Figure 4**). Taken together, these results indicate that *Ezh2* deletion in B cells negatively affected immature B cell differentiation in the BM. An increase in Pro-B cells may reflect a negative feedback loop in the BM to generate more functional B cells.

In the spleen, CD19^+^B220^+^ B cells were significantly reduced with EZH2 deficiency (**Figure 4A, gating strategies are in Supplementary Figure 5 and Supplementary Figure 6**). Proportions of transitional B cells (T2 B cells, **Figure 4B**), MZ B cells (**Figure 4C**), FO B cells (**Figure 4D**) and GC B cells (**Figure 4E**) were not significantly changed in CD19-Cre *Ezh2*^*fl/fl*^ compared to controls. Higher percentages of memory B cells were found in CD19-Cre *Ezh2*^*fl/fl*^ mice (**Figure 4F**). Differentiation of plasmablasts (**Figure 4G**) was inhibited in CD19-Cre *Ezh2*^*fl/fl*^ compared to controls. We found sex differences in B cell development in the spleen. CD19^+^B220^+^ B cells, frequency of T2 B cells, and GC B cells were significantly decreased in male CD19-Cre *Ezh2*^*fl/fl*^ mice compared to littermate controls, but not in females (**Supplementary Figure 7**), which might relate to the inherent difference that male controls had higher percentage of splenic B cells compared to female *Ezh2*^*fl/fl*^ mice (**Supplementary Figure 7A**). These results indicate that B cell differentiation in the spleen was disrupted with EZH2 deficiency. ASCs were significantly reduced, which might explain lower anti-dsDNA antibody production in CD19-Cre *Ezh2*^*fl/fl*^ mice.

**Figure 4.**
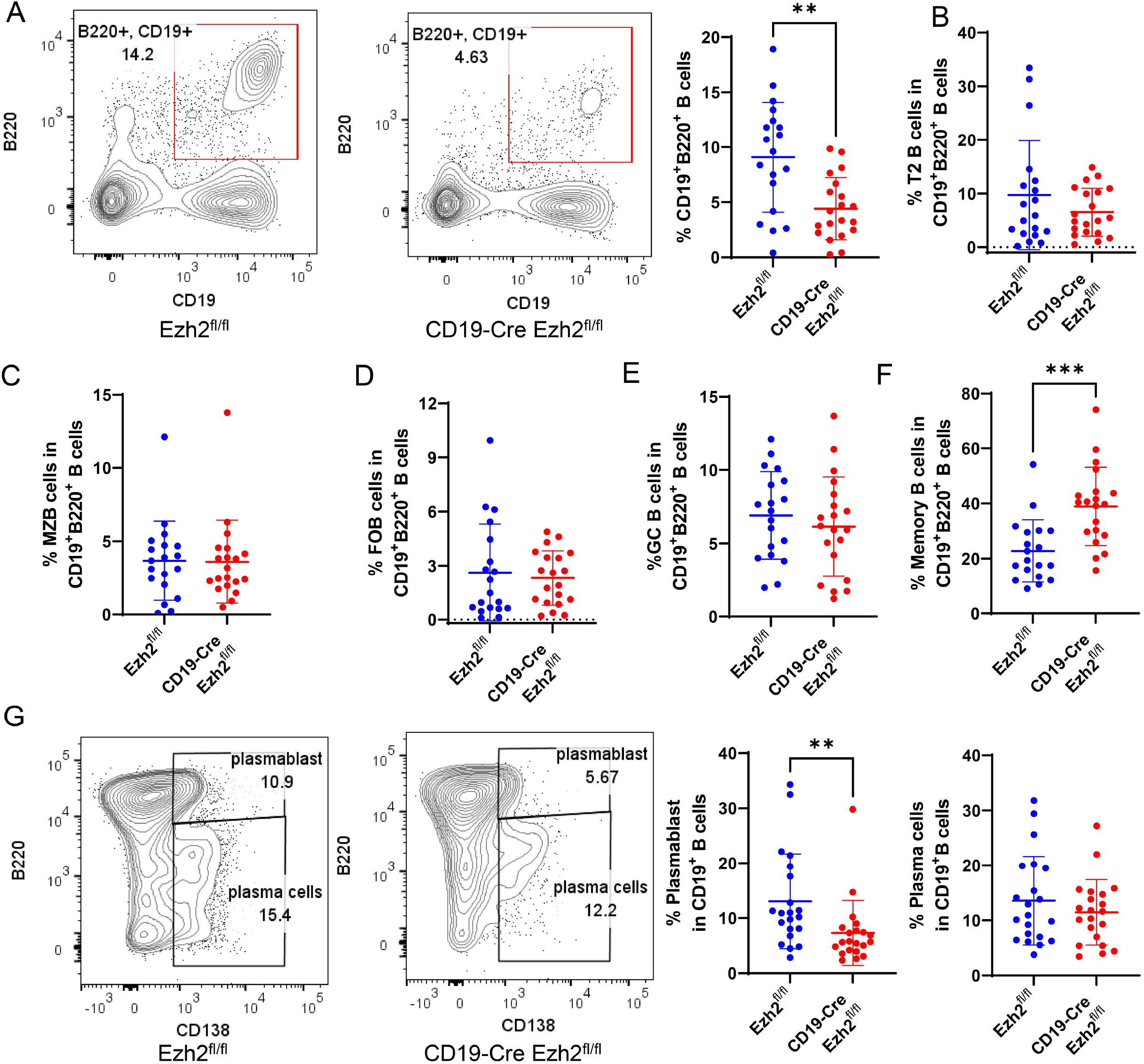
EZH2 deficiency in splenic B cells inhibits antibody secreting cell differentiation. **(A)** Representative flow cytometry plots, and summary data of splenic CD19^+^B220^+^ B cells in *Ezh2*^*fl/fl*^ (n=19) and CD19-Cre *Ezh2*^*fl/fl*^ (n=20) mice. **(B)** Proportion of transitional B cells (T2), **(C)** marginal zone B cells (MZB), **(D)** follicular B cells (FOB), **(E)** geminal center B cells (GC B) and **(F)** memory B cells in CD19^+^B220^+^ B cells in *Ezh2*^*fl/fl*^ (n=19) and CD19-Cre *Ezh2*^*fl/fl*^ (n=20) mice. **(G)** Representative flow cytometry plots and frequency data of, CD19^+^B220^high^CD138^+^ plasmablast and CD19^+^B220^low-medium^CD138^+^ plasma cellsin CD19^+^B220^+^ B cells in *Ezh2*^*fl/fl*^ (n=19) and CD19-Cre *Ezh2*^*fl/fl*^ (n=20) mice. Data are shown as mean ± SD. ** *p*<0.01, \*\*\**p*<0.001, two-tailed Mann-Whitney test.

### Single cell RNA-seq reveals B cell subset compositional and functional changes with Ezh2 deletion

To explore why EZH2 deficiency in B cells inhibited ASC differentiation, single cell RNA-sequencing was performed in FACS-sorted splenic CD19^+^ B cells. In total, 21 splenic cell populations were clustered based on cell transcriptional profile. Cell identity of each cluster was determined by evaluating differentially expressed genes within the clusters (**Supplementary Table 1**). We annotated each cluster and combined two GC B cell clusters, six plasmablast clusters, and 5 plasma cell clusters into single clusters. Twelve final cell clusters were retained (**Figure 5A**), 4 of which were non-B cell clusters which were excluded from subsequent analysis (clusters 7, 9, 10, and 12). Marker genes defining each B cell cluster are presented in **Figure 5B**. Signature genes for pre-memory B cells (cluster 1) included *Bach2* and *Ptprc* [29], and genes essential in B cell development including *Ebf1, Malt1, and Pag1*. Marker genes of MZ B cells (cluster 2) included high expression of MHC II genes (*H2-Eb1, H2-Aa, Cd74*) and *Fcmr* (encodes a Fc receptor of IgM) [30]. FO B cells (cluster 3) are characterized by high expression of *Ly6d [31], Cd79a, Ighd, Fcer2a* [29], *Ms4a1*, and *Cd79b*. ASCs including plasma blasts and plasma cells highly expressed *Xbp1 and Sdc1[32]*. It has been reported that *Prdm1* is highly induced in plasma cells but not plasmablasts [33], based on which we distinguished plasmablasts with low *Prdm1* expression and high levels of *Jchain* and *Ssr4* (cluster 4) from plasma cells with high expression levels of *Prdm1* and the one of its target genes *Trp53inp1* (cluster 6). GC B cells (cluster 5) were characterized by expression of *Tubb5, Mki67, Top2a* [29], *Pclaf* [34], *and Cenpf* [35]. Memory B cells (cluster 8) overexpressed *Zeb2, Cd38* [31], *Cd80* [36], *Zbtb20* [32], and *Sox5*[37]. A cluster (cluster 11) characterized by genes important for Pro-B cells and Pre-B cells including *Vpreb3, Igll1, Sox4, Vpreb1, Il7r, Cd93*, and *Rag1* was also in spleen cells.

**Figure 5.**
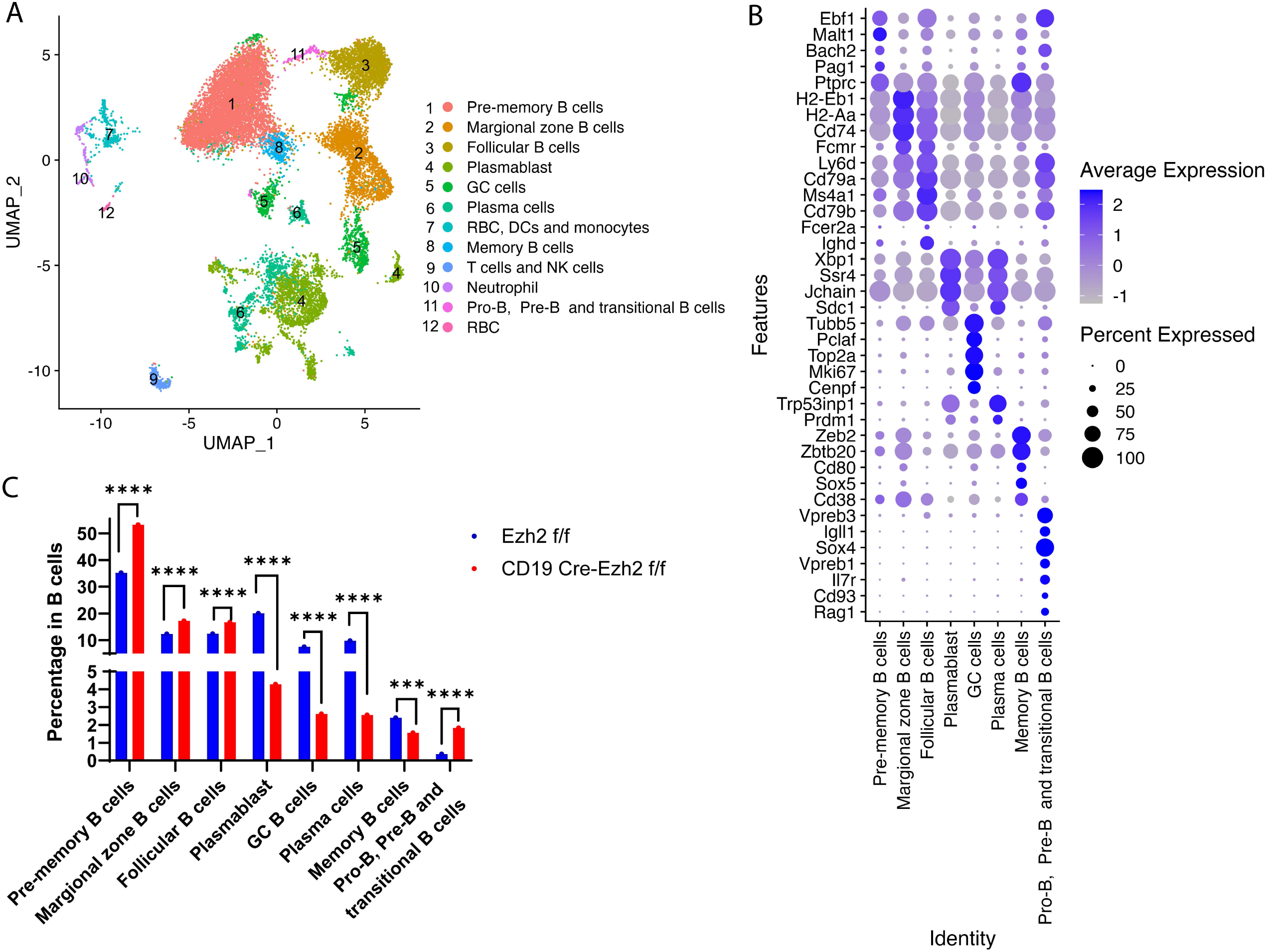
Single-cell RNA sequencing of female splenic B cells from *Ezh2*^*fl/fl*^ (n=2) and CD19-Cre *Ezh2*^*fl/fl*^ (n=2) mice. **(A)** UMAP plot of B cells and other cell types combining *Ezh2*^*fl/fl*^ and CD19-Cre *Ezh2*^*fl/fl*^ mice. **(B)** Marker genes of defining B cell clusters using single-cell RNA sequencing data from *Ezh2*^*fl/fl*^mice. **(C)** Frequencies of B cell identities in *Ezh2*^*fl/fl*^ and CD19-Cre *Ezh2*^*fl/fl*^ mice. \*\*\**p*<0.001, \*\*\*\**p*<0.0001, Fisher’s exact test of normalized cell frequencies in each B cell subtype. GC, germinal center; RBC, red blood cells; DCs, dendritic cells; NK cells, natural killer cells.

We analyzed the proportions of each cell cluster to determine differences in B cell subset composition after excluding non-B cell clusters between EZH2-deficient and control mice. Transitional B cells, MZ B cells and FO B cells were significantly expanded in CD19-Cre *Ezh2*^*fl/fl*^ compared to *Ezh2*^*fl/fl*^ mice (**Figure 5C**). Further, scRNA-seq results showed reduced differentiation of GC B cells, plasmablasts, and plasma cells in CD19-Cre *Ezh2*^*fl/fl*^ mice, confirming our flow cytometry data (**Figure 5C**). Pre-memory B cells were significantly expanded, while memory B cells were reduced in CD19-Cre *Ezh2*^*fl/fl*^ mice compared to controls.

### XBP1 as a downstream target of EZH2 in B cells regulating plasmablast/plasma cell differentiation

Because we observed reduced anti-dsDNA antibody production with B cell-*Ezh2* deletion, and reduced plasmablasts/plasma cells (ASCs), we hypothesized that reduced ASC differentiation is probably key in explaining disease amelioration in our mouse model. To understand why ASC differentiation was defective with EZH2 deficiency in lupus mouse B cells, and because T cell-dependent and T cell-independent ASCs primarily arise from GC and MZ B cells [38], we examined differentially expressed genes (DEGs) in GC and MZ B cell subsets in CD19-Cre *Ezh2*^*fl/fl*^ mice compared to *Ezh2*^*fl/fl*^ control mice (Complete lists of DEGs in all clusters are shown in **Supplementary Table 2**). There were 322 and 371 significant DEGs in GC B and MZ B cell subsets, respectively, with 73 common differentially expressed genes in both GC and MZ B cell subsets (**Figure 6A**). Functional enrichment analysis of these 73 DEGs (50/73 were used for analysis, the other 23 were mouse immunoglobulin heavy and light chain genes) was performed using ToppGene (https://toppgene.cchmc.org/enrichment.jsp), and revealed one Gene Ontology related to B cell activation (GO: 0042113, **Supplementary Table 3**), which included 7 genes: *MZB1, BATF, IGHG1, XBP1, CD79A, CD81*, and *LGALS1* (**Table 1**). The expression of XBP1, which is a key regulator for B cell development and plasmablast/plasma cell differentiation was significantly downregulated with *Ezh2* deletion (P= 1.4×10^−3^ and 7.2×10^−6^, in GC and MZ B cells, respectively). The expression of BACH2, which has been previously shown to regulate XBP1 expression in B cells [18], was not different between EZH2-deficient and control mice (not shown), suggesting the involvement of alternative mechanisms for XBP1 regulation in our disease model.

**Figure 6.**
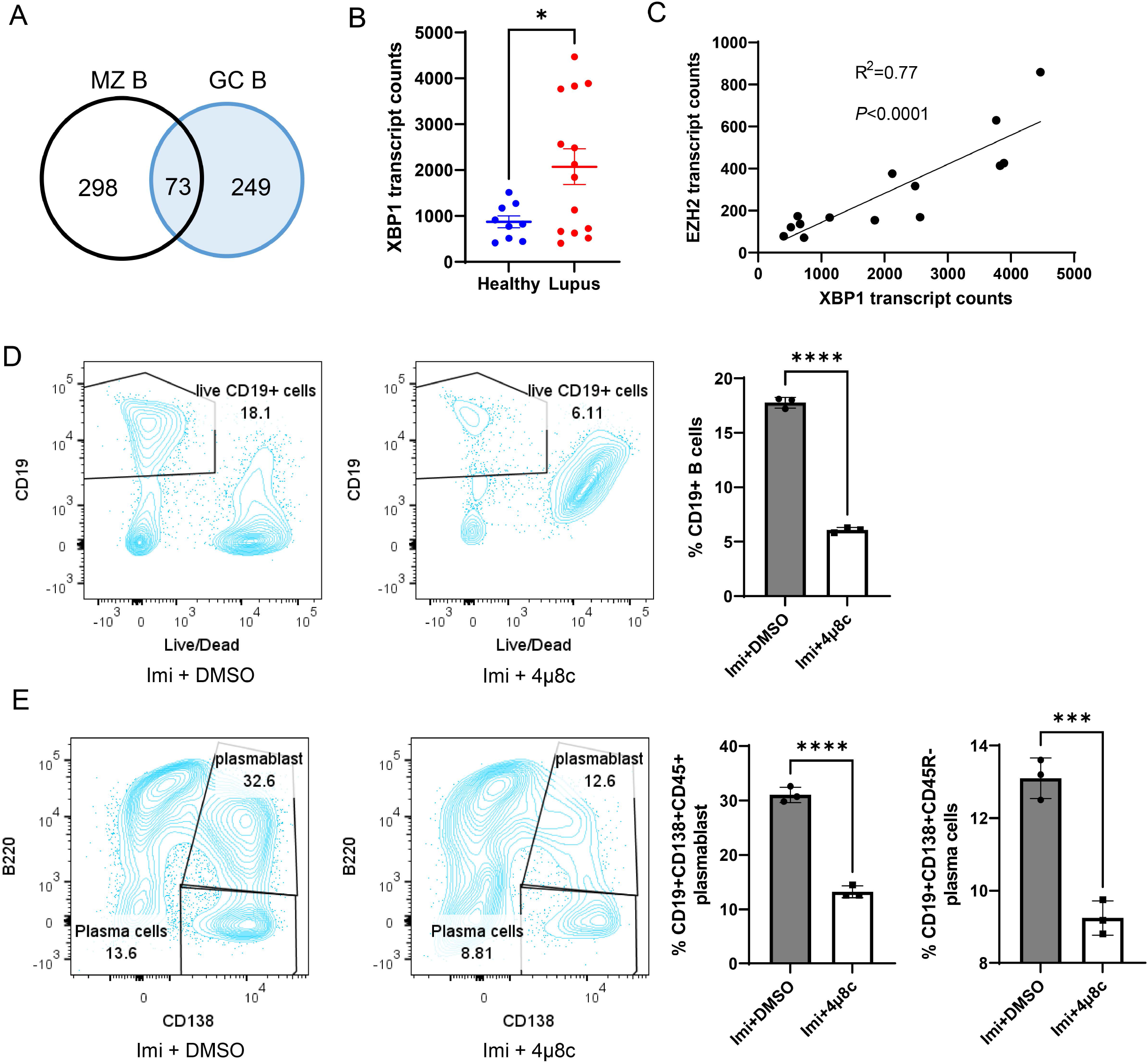
XBP1 expression in human CD19^+^ cells and XBP1 inhibition in B cells from MRL/*lpr* mice. **(A)** Venn diagrams depicting the number of differentially expressed genes in marginal zone (MZ) B cells and germinal center (GC) B cells between *Ezh2*^*fl/fl*^ and CD19-Cre *Ezh2*^*fl/fl*^ mice in single-cell RNA sequencing. **(B)** XBP1 transcript counts in CD19^+^ B cells from female healthy controls (n=9) and female lupus patients (n=14) in RNA-sequencing. Data are presented as mean ± SE. * *p*<0.05, unpaired two-tailed *t* test. **(C)** Correlation plot of XBP1 and EZH2 mRNA expression levels from female lupus patients (n=14). Representative flow cytometry plots and summary data of **(D)** CD19^+^ B cells, and **(E)** plasmablasts and plasma cells isolated from the spleen of MRL/*lpr* mice after stimulation with 3 μg/ml TLR7 agonist imiquimod (Imi) with and without 20 μM XBP1 inhibitor 4μ8c for 3 days. DMSO was used as vehicle control. Results are presented as mean ± SD, data are representative of 3 independent experiments. \*\*\**p*<0.001, \*\*\*\**p*<0.0001, unpaired 2-tailed *t* test.

**Table 1.**
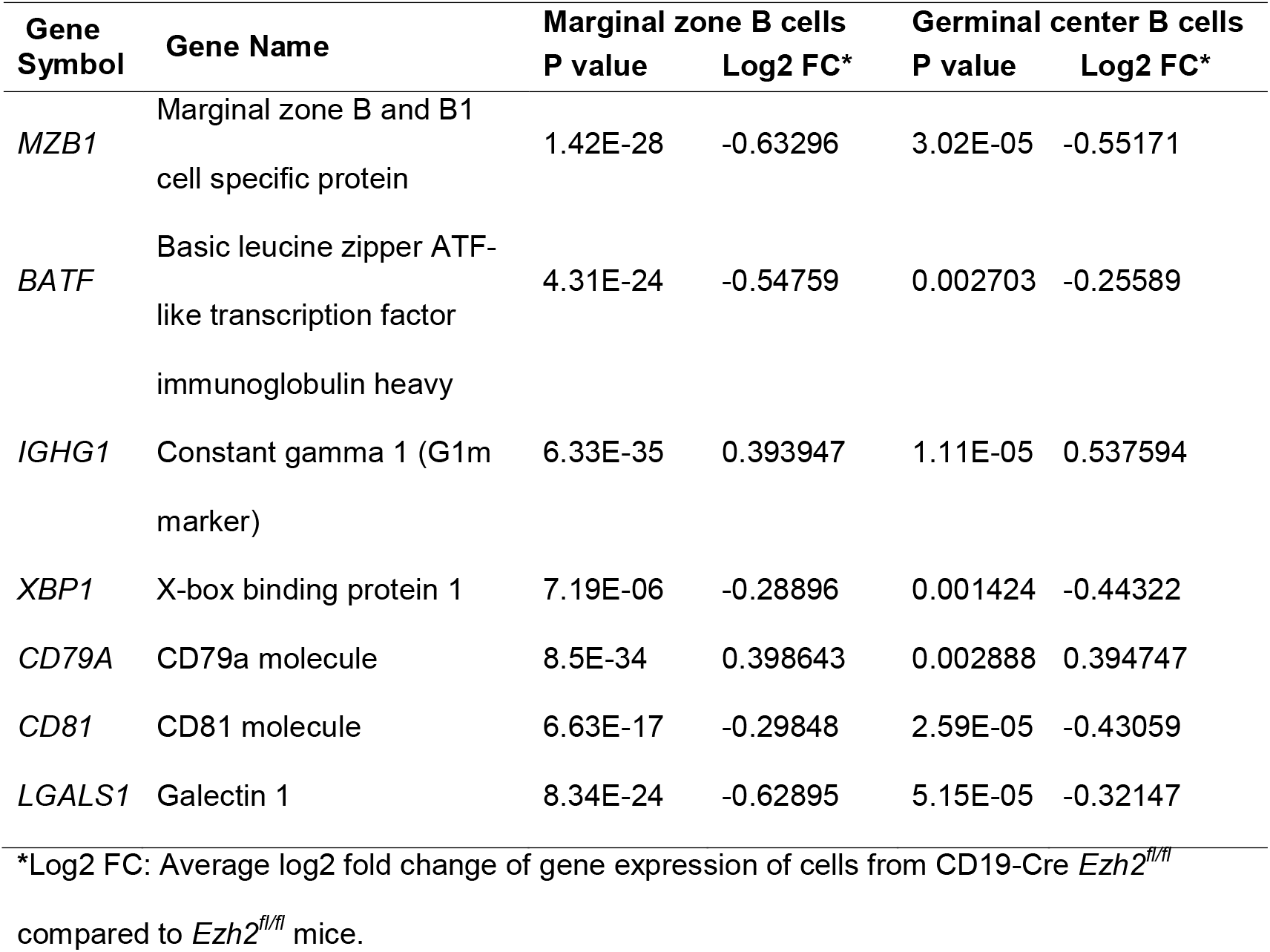
Differentially expressed genes of B cell activation in marginal zone B cells and germinal center B cells between *Ezh2*^*fl/fl*^ and CD19-Cre *Ezh2*^*fl/fl*^ mice.

Analyzing mRNA expression levels in peripheral blood B cells, comparing lupus patients with normal healthy controls, revealed significant increase in XBP1 expression in lupus patients (**Figure 6B**). Indeed, there was a strong correlation between EZH2 expression and XBP1 expression levels in CD19^+^ B cells from lupus patients (R^2^=0.77, **Figure 6C**).

To determine if decreased XBP1 leads to reduced plasmablast and plasma cell differentiation in MRL/*lpr* mice, we isolated B cells from the spleen of wild-type MRL/*lpr* mice and stimulated them *in vitro* with imiquimod to induce plasmablast differentiation in the presence or absence of the XBP1 inhibitor 4μ8c. XBP1 inhibition resulted in significant suppression of imiquimod-induced B cell expansion (**Figure 6D**), and defective plasmablast/plasma cell differentiation in MRL/*lpr* B cells, similar to B cell specific EZH2-deficient mice (**Figure 6E**). These data suggest that XBP1 downregulation in EZH2-deficient B cells might explain reduced plasmablast and plasma cell differentiation in CD19-Cre *Ezh2*^*fl/fl*^ mice.

### Class switch recombination (CSR) is defective in B cell EZH2-deficient mice

Upon T cells priming, FO B cells will undergo an intrachromosomal DNA recombination process to change the immunoglobulin (Ig) heavy chain constant region, which is called class switch recombination (CSR), to better protect against pathogens. IgM or IgD isotypes can be replaced by IgG, IgA, or IgE through CSR without altering antigen specificity [39]. Dysregulation of CSR can lead to autoimmunity [40]. We performed single cell BCR sequencing, and integrated the results with scRNA-seq-defined clusters to investigate whether CSR was affected with EZH2 deficiency in B cells. We found different Ig heavy chain and light chain isotypes distribution across the two mouse groups (**Figure 7A**). Specifically, *IGHM* (immunoglobulin heavy constant mu) which defines the IgM isotype was significantly higher, and *IGHG*s (immunoglobulin heavy constant gamma) including *IGHG1, IGHG2C*, and *IGHG3* which encode IgG1, IgG2c, and IgG3, respectively, showed a trend to be lower in the plasma cells from CD19 Cre-*Ezh2*^*fl/fl*^ mice compared to *Ezh2*^*fl/fl*^ control mice (**Figure 7B**). This implies that *IGHM* switching to *IGHGs* was impaired in the GC. These results suggest that EZH2 deficiency inhibits IGHM class switch recombination (CSR), resulting in accumulation of IGHM and decreased IGHG isotypes in MRL/*lpr* mice.

**Figure 7.**
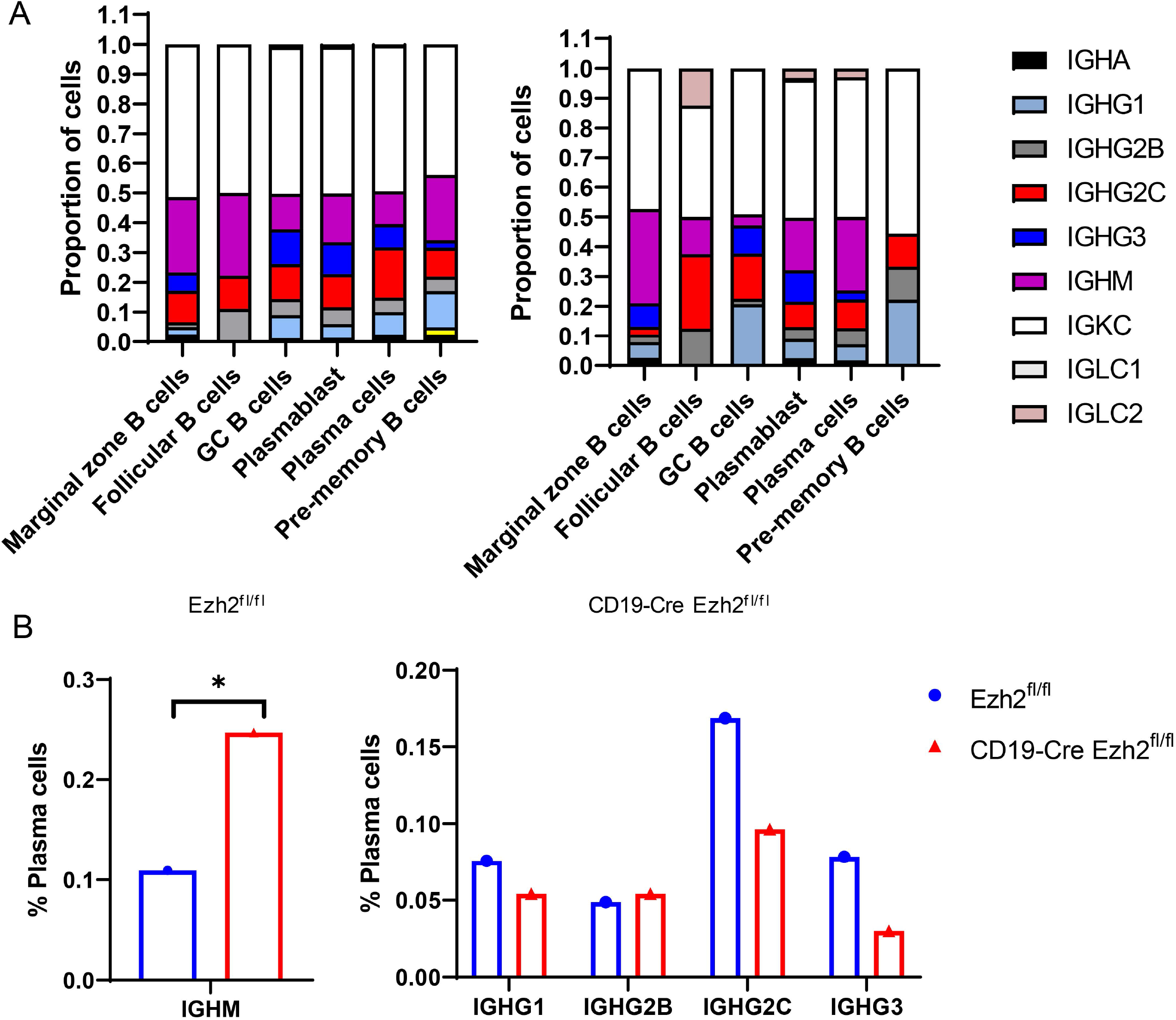
Single cell B cell receptor (BCR) sequencing of splenic B cells from *Ezh2*^*fl/fl*^ and CD19-Cre *Ezh2*^*fl/fl*^ mice. **(A)** BCR isotype distribution of 6 B cell clusters (n=6414 cells and n=672 cells respectively in *Ezh2*^*fl/fl*^ and CD19-Cre *Ezh2*^*fl/fl*^ mice). (**B**) Proportion of plasma cells with IGHM and IGHG isotypes (plasma cell n=1518 in *Ezh2*^*fl/fl*^ mice and n=166 in CD19-Cre *Ezh2*^*fl/fl*^). * *p*<0.05, Fisher’s exact test of normalized cell frequencies in plasma cells.

## Discussion

We found that MRL/*lpr* mice with EZH2 deficiency in B cells developed less severe lupus-like phenotypes, including lowered levels of serum autoantibodies and proteinuria, and improved glomerulonephritis. B cell differentiation into ASCs was inhibited with EZH2 deficiency, possibly as a result of XBP1 downregulation. In addition, class switch recombination appears to be defective in MRL/*lpr* mice with EZH2 deficiency in B cells.

B cells play an important role in the pathogenesis of autoimmune diseases, including lupus. B cell targeting therapies, such as BAFF inhibitors, are successfully used in the treatment of lupus and lupus-related renal involvement [41]. EZH2 is a key epigenetic regulator that is critically involved in B cells development and differentiation [42]. We found that EZH2 deficiency significantly decreased total splenic B cell populations (**Figure 4A and Supplementary Figure 7A**), and inhibited the differentiation of ASCs (**Figure 4G and Supplementary Figure 7G**). These data provided supporting evidence that EZH2 is indispensable for peripheral B cell development. At the same time, memory B cells were significantly increased in EZH2-deficient MRL*/lpr* lupus-prone mice (**Figure 4F and Supplementary Figure 7F**). In NP-CGG_25-36_ (4-Hydroxy-3-nitrophenylacetyl-Chicken Gamma Globulin) immunized mice models, both of GC B cells and memory B cells formation were impaired by EZH2 inactivation [43], which is contrary to what we observed in lupus-like MRL/*lpr* mice. This discrepancy may be because antigen exposure is consistent in lupus-prone MRL/*lpr* mice compared to one-time immunization models. Our data suggest that in MRL/*lpr* mice, EZH2 deficiency possibly results in inefficient differentiation of memory B cells into GC B cells and plasmablasts/plasma cells, explaining accumulation and therefore increased numbers of memory B cells. Furthermore, some memory B cells in the CD19-Cre *Ezh2*^*fl/fl*^ model may arise from activated B cells in the marginal zones [44], which were not reduced by EZH2 deficiency in our model (**Figure 4C, Supplementary Figure 7C and Figure 5C**).

Our data demonstrated downregulation of XBP1 in EZH2-deficient B cells, which is a key transcription factor that mediates B cell differentiation [45]. This suggest that the effect of EZH2 deficiency upon the differentiation of B cells could be mediated through XBP1 reduction. Indeed, we demonstrate similar effects on B cell differentiation in MRL/*lpr* mice, *ex vivo*, with an XBP1 inhibitor. We also demonstrate a strong positive correlation between mRNA expression levels of EZH2 and XBP1 in human lupus B cells. How EZH2 regulates transcription factor XBP1 needs more investigation. Zhang et al. showed that EZH2 epigenetically suppresses BACH2 expression in human CD19^+^ cells from healthy donors, resulting in increased expression of downstream PRDM1 and XBP1 [18]. In our scRNA-seq data, the expression of BACH2 in B cell clusters was not affected by *Ezh2* deletion (**Supplementary Table 2**). Therefore, our data suggest that XBP1 might be regulated by EZH2 through other mechanisms independent of the BACH2/PRDM1 axis.

CSR happens before GC formation and also in the GC to switch IgD and IgM to IgG, IgA, and IgE isotypes [46]. Our sc-BCR sequencing suggests impaired ability of *IGHM* to switch to *IGHGs* in the plasma cells from CD19-Cre *Ezh2*^*fl/fl*^ mice (**Figure 7B**). Previous studies showed that IgG anti-dsDNA was positively correlated with glomerulonephritis in SLE patients, while IgM anti-dsDNA was protective from lupus nephritis in mouse models and negatively correlated with human glomerulonephritis [47, 48]. This is consistent with our results in CD19-Cre *Ezh2*^*fl/fl*^ mice (**Figure 2B-2D**).

In conclusion, EZH2 deficiency in B cells ameliorated lupus-like disease in MRL/*lpr* mice. EZH2 deficiency impeded B cell development and differentiation in the bone marrow and spleen. B cell *Ezh2* deletion led to significant reduction in ASCs. Single cell transcriptomics and *in vitro* data showed that XBP1 is a downstream target of EZH2 in MRL/*lpr* mice and might at least in part mediate the effects of EZH2 deficiency upon B cell development. CSR was also defective with EZH2 deficiency in B cells, possibly impairing pathogenic autoantibody production. Our results provided mechanistic evidence supporting targeting EZH2 as a potential novel therapeutic option for SLE.

## Contributorship

- All authors fulfilled the following criteria:
- Substantial contributions to the conception or design of the work, or the acquisition, analysis or interpretation of data.
- Drafting the work or revising it critically for important intellectual content.
- Final approval of the version published.

## Acknowledgements

This work was supported by the National Institute of Allergy and Infectious Diseases (NIAID) of the National Institutes of Health (NIH) grant number R01 AI097134. We are grateful to Dr. Mark Shlomchik for providing CD19-Cre MRL/*lpr* mice for our studies. We are also grateful to Dr. Sebastien Gingras of the Innovative Technologies Development Core (Department of Immunology, University of Pittsburgh) for his help with the design and identification of the *Ezh2* floxed mouse on MRL/*lpr* background, as well as Dr. Chunming Bi and Zhaohui Kou of the Mouse Embryo Services Core (Department of Immunology, University of Pittsburgh) for microinjection of zygotes and production of the mice. We thank Joshua Michel and Valerie Miller in the Rangos Research Flow Core for their help with flow cytometry data analysis and sorting. We thank Tracy Tabib of the Single Cell Core for preparing single cell sequencing library and help with data analysis. We thank Michele Mulkeen of the histology core for her help with tissue sectioning and staining. We also thank Luis Espinoza and Mckenna M. Bowes for helping to collect mouse tissues.

## Ethical approval information

The study was approved by the Institutional Animal Care and Use Committee (IACUC) of the University of Pittsburgh.

## Data sharing statement

All data are included in the manuscript or supplementary material.

## Patient and public involvement

It was not appropriate or possible to involve patients or the public in the design, or conduct, or reporting, or dissemination plans of our research.

