## Supplementary Materials and Figures for "*Ezh2* knockout in B cells impairs plasmablast differentiation and ameliorates lupus-like disease in MRL/*lpr* mice"

**Supplementary Material**

**Supplementary Methods**

*Mouse genotyping method*. DNA from mice tail (2-5 mm) was isolated following the instruction of genomic DNA isolation kit (Lamda Biotech, St. Louis, MO, USA). Genotyping primers were as follows: *Ezh2* F1: GAACAAGGGGTTCGGGATCA; *Ezh2* R1: AGCACACCCACTTACACAGG. *CD19* forward primer: AATGTTGTGCTGCCATGCCTC; *CD19-Cre* reverse primer: TTCTTGCGAACCTCATCAC; *CD19* reverse primer: GTCTGAAGCATTCCACCGGAA. Q5 high-fidelity DNA polymerase (NEW ENGLAND BioLabs) was used for genotyping. PCR components and the PCR reaction used are shown in Table 1 and Table 2 below. PCR products were run on a 2% agarose gel. *CD19* wild-type (WT) shows a band at 550 bp and *CD19-Cre* shows a band at 220 bp. *Ezh2* PCR product is then digested with EcoRI-HF restriction enzyme (NEW ENGLAND BioLabs) as shown in Table 3 to interpret genotyping. After digestion, *Ezh2* WT shows a band at 1075 bp and *Ezh2^fl/fl^* shows three bands at 491 bp, 421 bp, and 243 bp on a 2% agarose gel.

Table 1. PCR components

| Component | 25 µl Reaction | |
| --- | --- | --- |
| 5X Q5 Reaction Buffer | | 5 |
| 10 mM dNTPs | | 0.5 |
| 10 µM Forward Primer | | 1.25 |
| 10 µM Reverse Primer | | 1.25 |
| Q5 High-Fidelity DNA Polymerase | | 0.25 |
| Nuclease-Free Water | | 15.75 |

Add 1 ul DNA isolated from tails

Table 2. PCR reaction

| Step | Temp | Time |
| --- | --- | --- |
| Initial Denaturation | 98°C | 30 seconds |
| 35 Cycles | 98°C | 10 seconds |
|  | 68°C | 20 seconds |
|  | 72°C | 30 seconds |
| Final Extension | 72°C | 2 minutes |
| Hold | 4–10°C |  |

Table 3. *Ezh2* PCR product Digestion

|  | | 1X (μl) |
| --- | --- | --- |
| *Ezh2* F1xR1 PCR Product | | 13.0 |
| 10X CutSmart Buffer | | 1.5 |
| EcoRI-HF (NEB 20,000 units/ml) | | 0.5 |
| Total volume | | 15.0 |
| Incubate at 37°C overnight |  | |

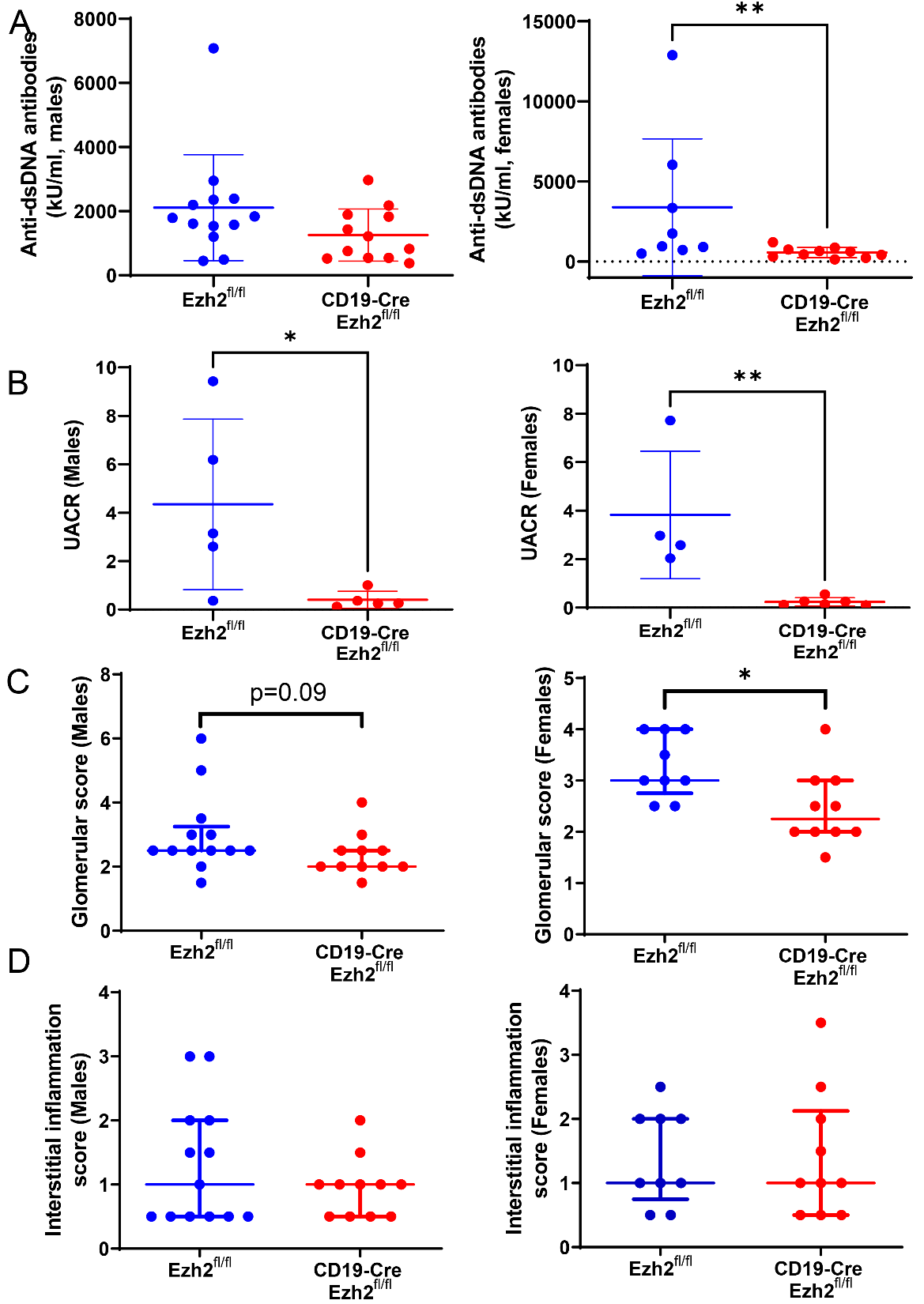
**Supplementary Figure 1. Sex-specific serum anti-dsDNA antibody levels and lupus nephritis data in *Ezh2^fl/fl^* and CD19-Cre *Ezh2^fl/fl^* MRL/*lpr* mice. (A)** Males (n=13 and n=12, respectively) and females (n=8 and n=10, respectively) anti-dsDNA antibody levels in *Ezh2^fl/fl^* and CD19-Cre *Ezh2^fl/fl^* mice. **(B)** Urine albumin to creatinine ratio (UACR) in males (n=5 and n=5, respectively) and females (n=4 and n=6, respectively) in *Ezh2^fl/fl^* and CD19-Cre *Ezh2^fl/fl^* mice. **(C)** Glomerular scores and **(D)** interstitial inflammation scores based on hematoxylin and eosin (H&E) stained kidney tissue in males (n=13 and n=11, respectively) and females (n=9 and n=10, respectively) in *Ezh2^fl/fl^* and CD19-Cre *Ezh2^fl/fl^* mice. Data in (**A**) and (**B**) are shown as mean ± SD. Data in (**C**) and (**D**) are presented as median with interquartile range, * *p*<0.05, ** *p*<0.01, two-tailed Mann-Whitney test.

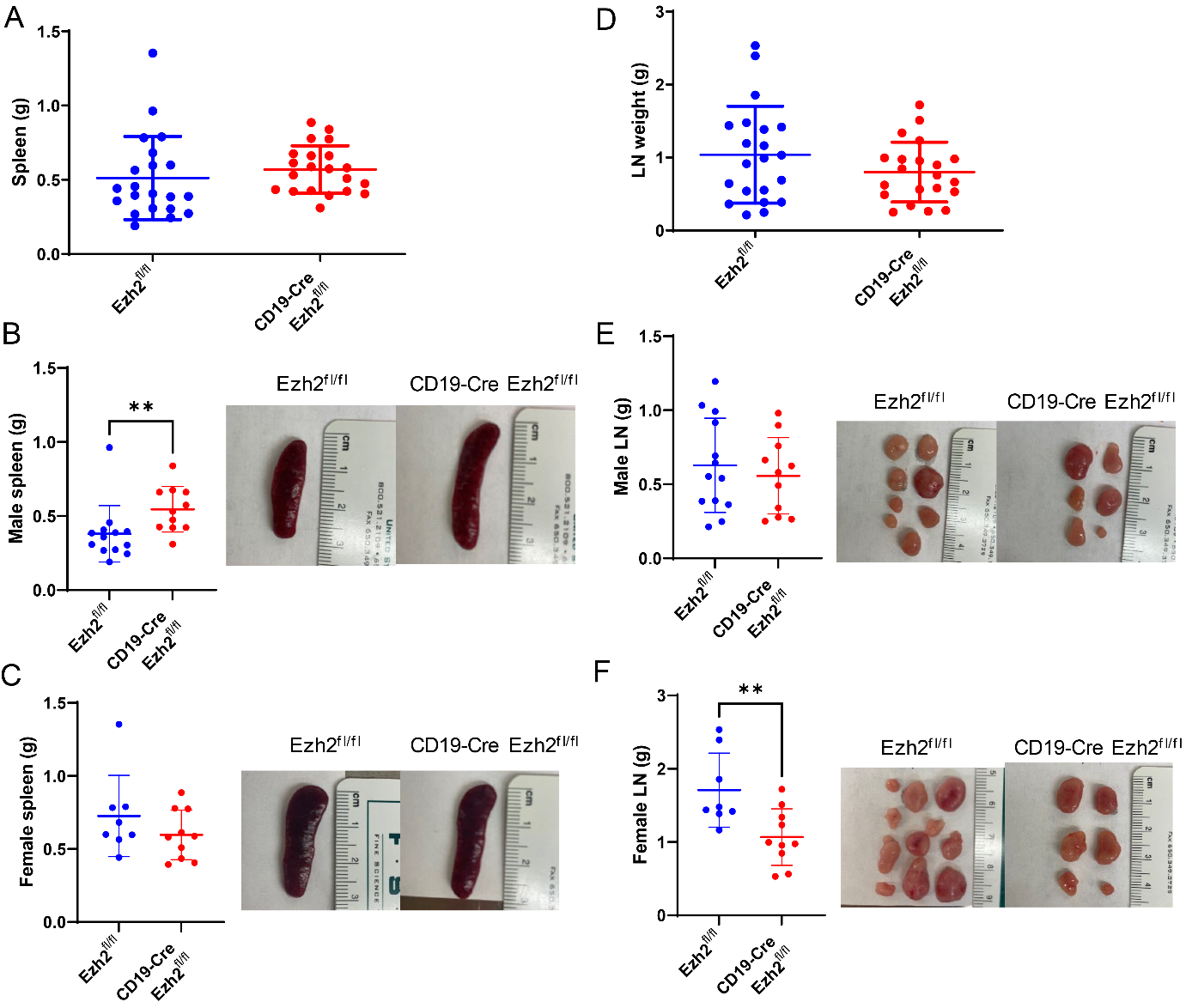

**Supplementary Figure 2. Splenomegaly and lymphadenopathy in CD19-Cre *Ezh2^fl/fl^* and *Ezh2^fl/fl^* MRL/*lpr* mice. (A)** Spleen weights in *Ezh2^fl/fl^* (n=21) and CD19-Cre *Ezh2^fl/fl^* (n=21) mice. Weight and representative images of the spleen in **(B)** male (n=13 and n=11, respectively) and **(C)** female (n=8 and n=10, respectively) in *Ezh2^fl/fl^* and CD19-Cre *Ezh2^fl/fl^* mice. **(D)** Weight of superficial cervical and deep cervical lymph nodes (LN) in *Ezh2^fl/fl^* (n=21) and CD19-Cre *Ezh2^fl/fl^* (n=21) mice. Weight and representative images of superficial cervical and deep cervical lymph nodes in **(E)** male (n=13 and n=11, respectively) and (**F**) female (n=8 and n=10, respectively) *Ezh2^fl/fl^* and CD19-Cre *Ezh2^fl/fl^* mice. Data are shown as mean ± SD, ** *p*<0.01, two-tailed Mann-Whitney test in (**B**) and unpaired 2-tailed *t* test in (**F**).

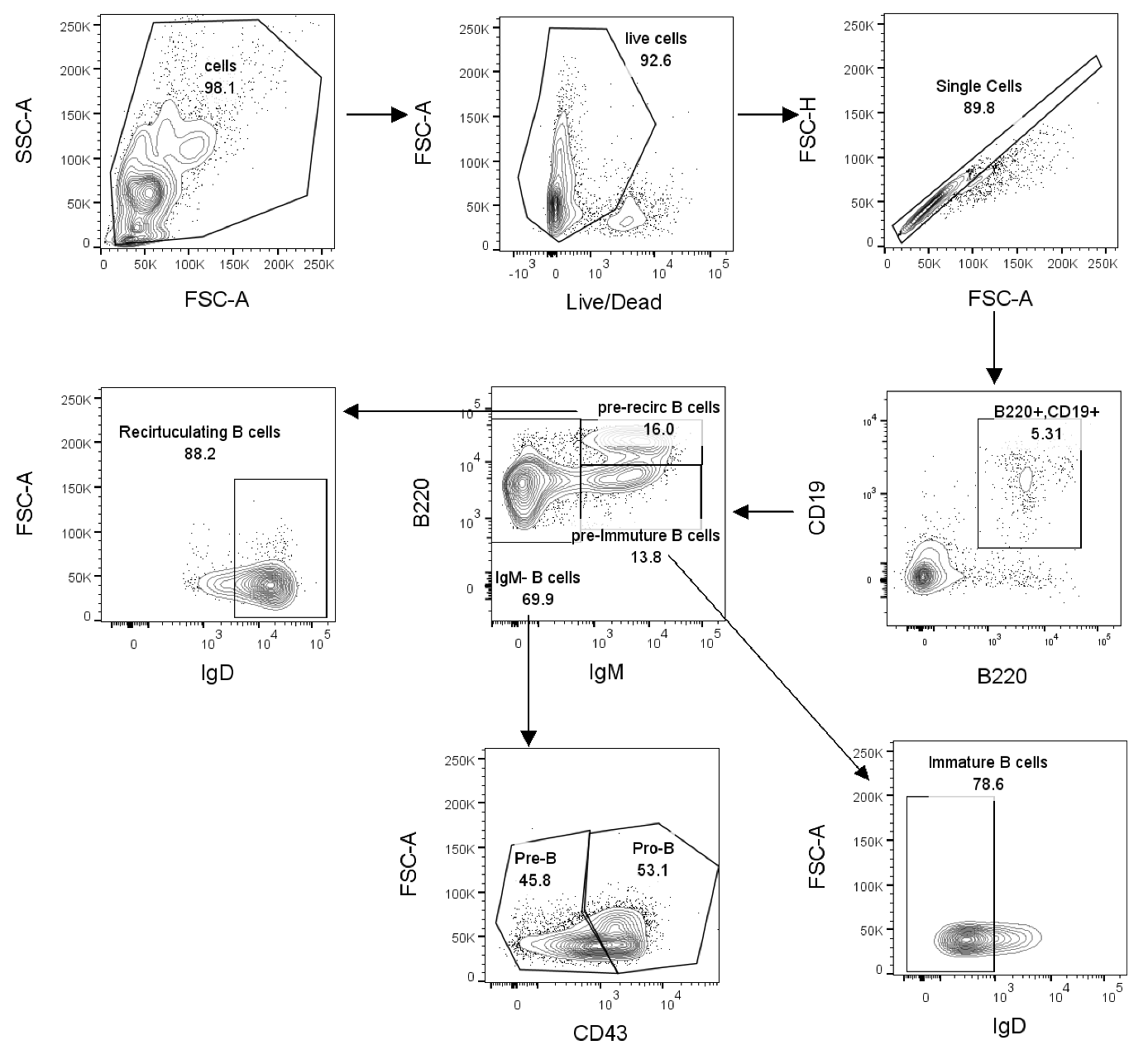

**Supplementary Figure 3. Gating strategy of B cells in the bone marrow.** Gating strategy of B cells (B220^+^CD19^+^), Pro-B cells (CD19^low^IgM^–^B220^+^CD43^+^), Pre-B cells (CD19^+^IgM^–^B220^+^CD43^–^), immature B cells (CD19^+^B220^int^IgM^+^IgD^–^), and recirculating B cells (CD19^+^B220^hi^IgM ^+^ IgD ^+^) in the bone marrow.

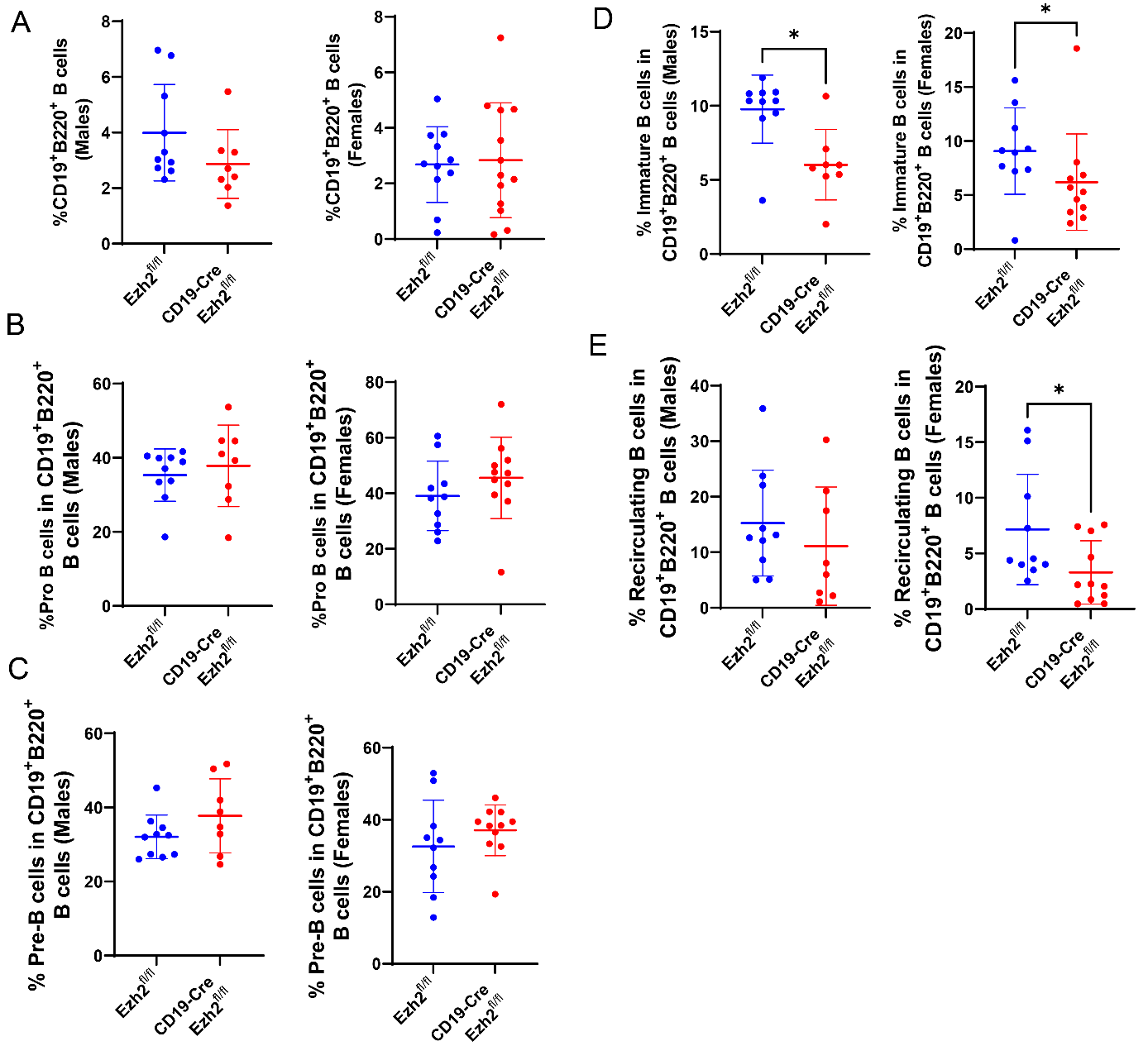

**Supplementary Figure 4. Sex-specific B cell development in the bone marrow of *Ezh2^fl/fl^* and CD19-Cre *Ezh2^fl/fl^* mice***.* Frequency of **(A)** CD19^+^B220^+^ B cells in male mice (n=10 and n=8, respectively) and female mice (n=11 and n=13, respectively), and proportion of **(B)** Pro-B cells, **(C)** Pre-B cells, **(D)** immature B cells, and **(E)** recirculating B cells in male mice (n=10 and n=8, respectively) and female mice (n=10 and n=11, respectively) in CD19^+^B220^+^ B cells in *Ezh2^fl/fl^* and CD19-Cre *Ezh2^fl/fl^* mice. * *p*<0.05, two-tailed Mann-Whitney test.

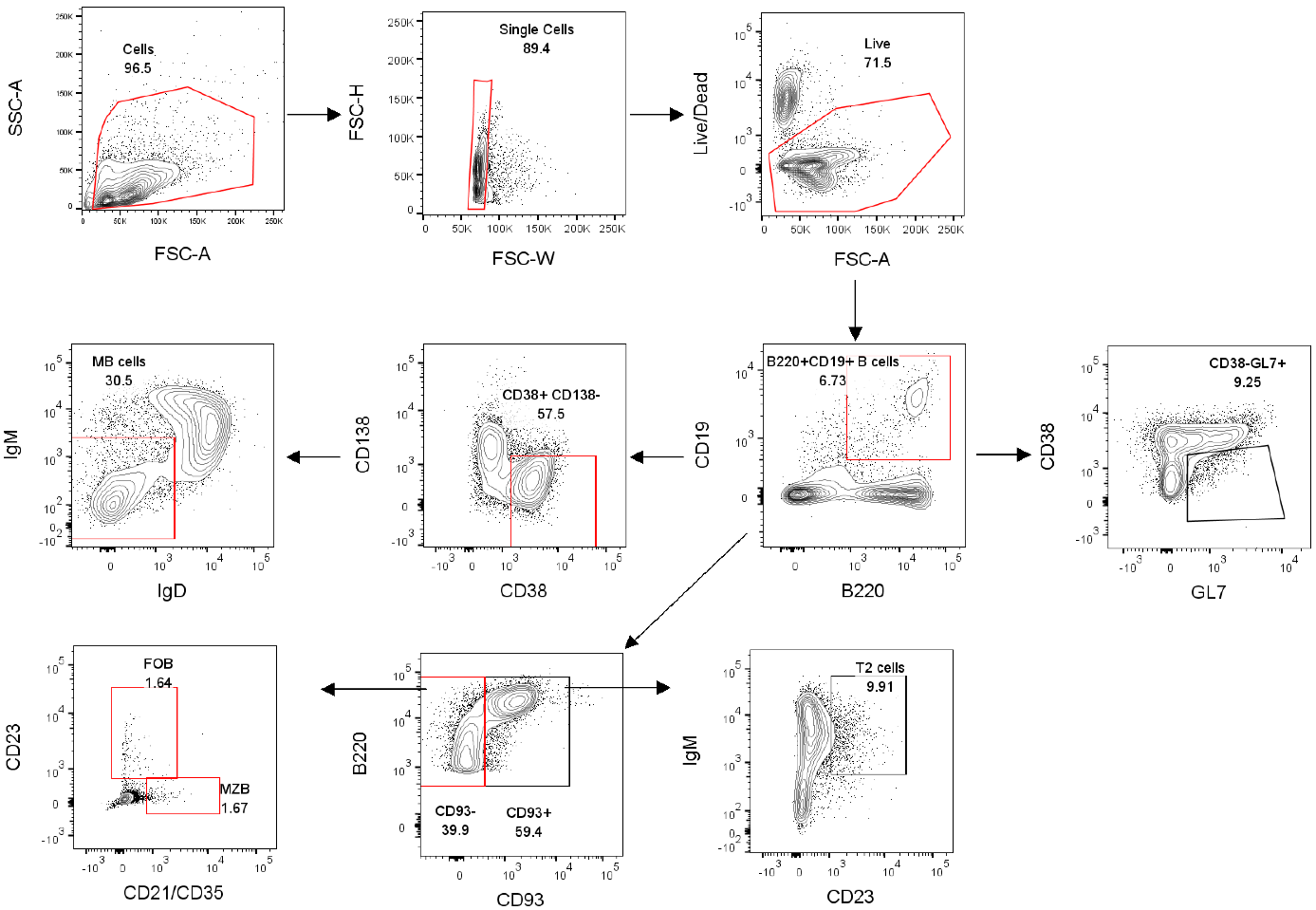

**Supplementary Figure 5. Gating strategy of B cells in the spleen.** Gating strategy of splenic B cells (B220^+^CD19^+^), transitional B cells (T2, CD19^+^B220^+^CD93^+^CD23^+^IgM^+^), marginal zone B cells (MZB, CD19^+^B220^+^CD93^-^CD21/CD35^+^CD23^-^), follicular B cells (FOB, CD19^+^B220^+^ CD93^-^CD21/CD35^neg-low^CD23^+^), germinal center B cells (CD19^+^B220^+^CD38^-^GL7^+^), and memory B cells (MB, CD19^+^ B220^+^CD138^-^CD38^+^IgM^-^IgD^-^).

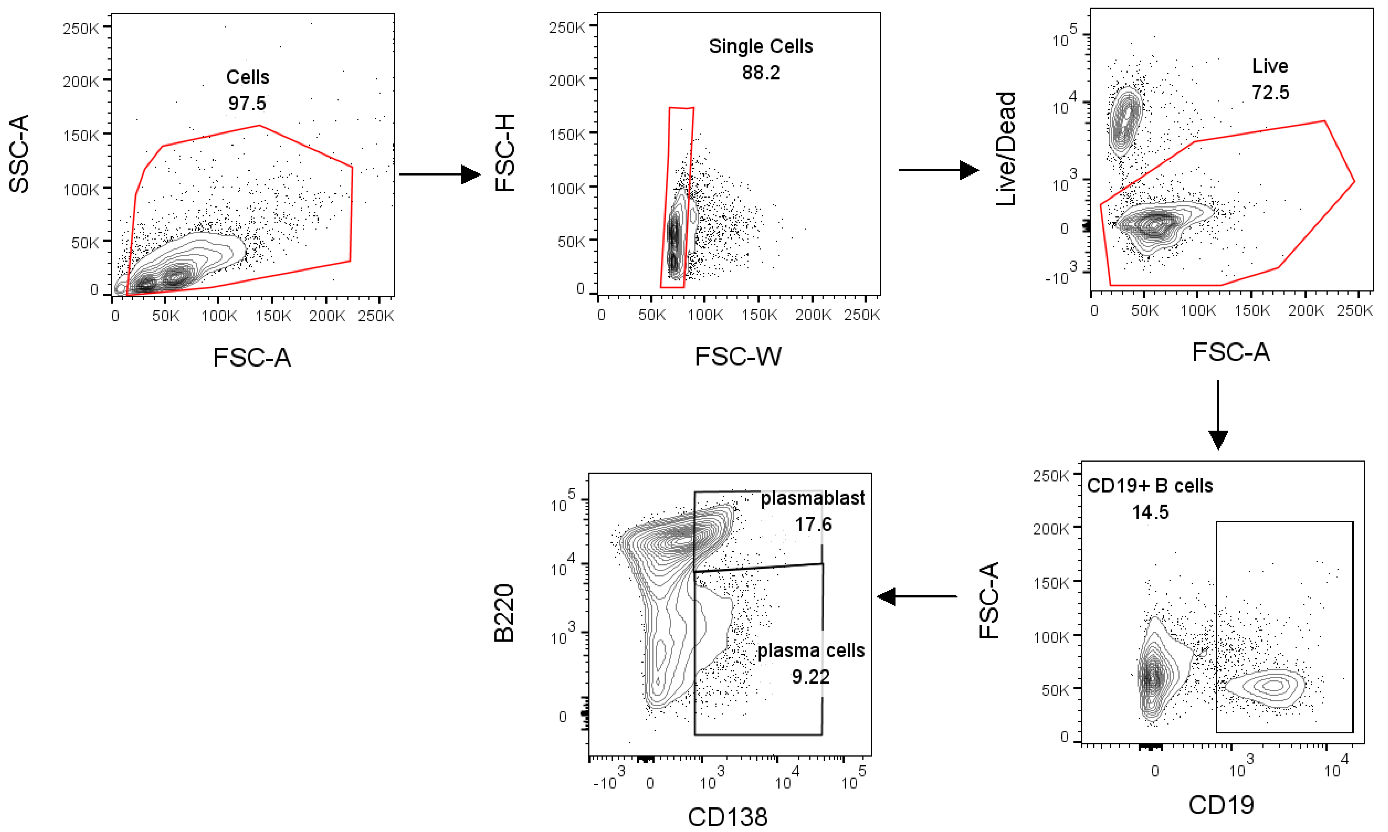

**Supplementary Figure 6. Gating strategy of plasma cells and plasmablasts in the spleen.** Gating strategies of plasmablasts (CD19^+^B220^high^CD138^+^) and plasma cells (CD19^+^B220^low-medium^CD138^+^).

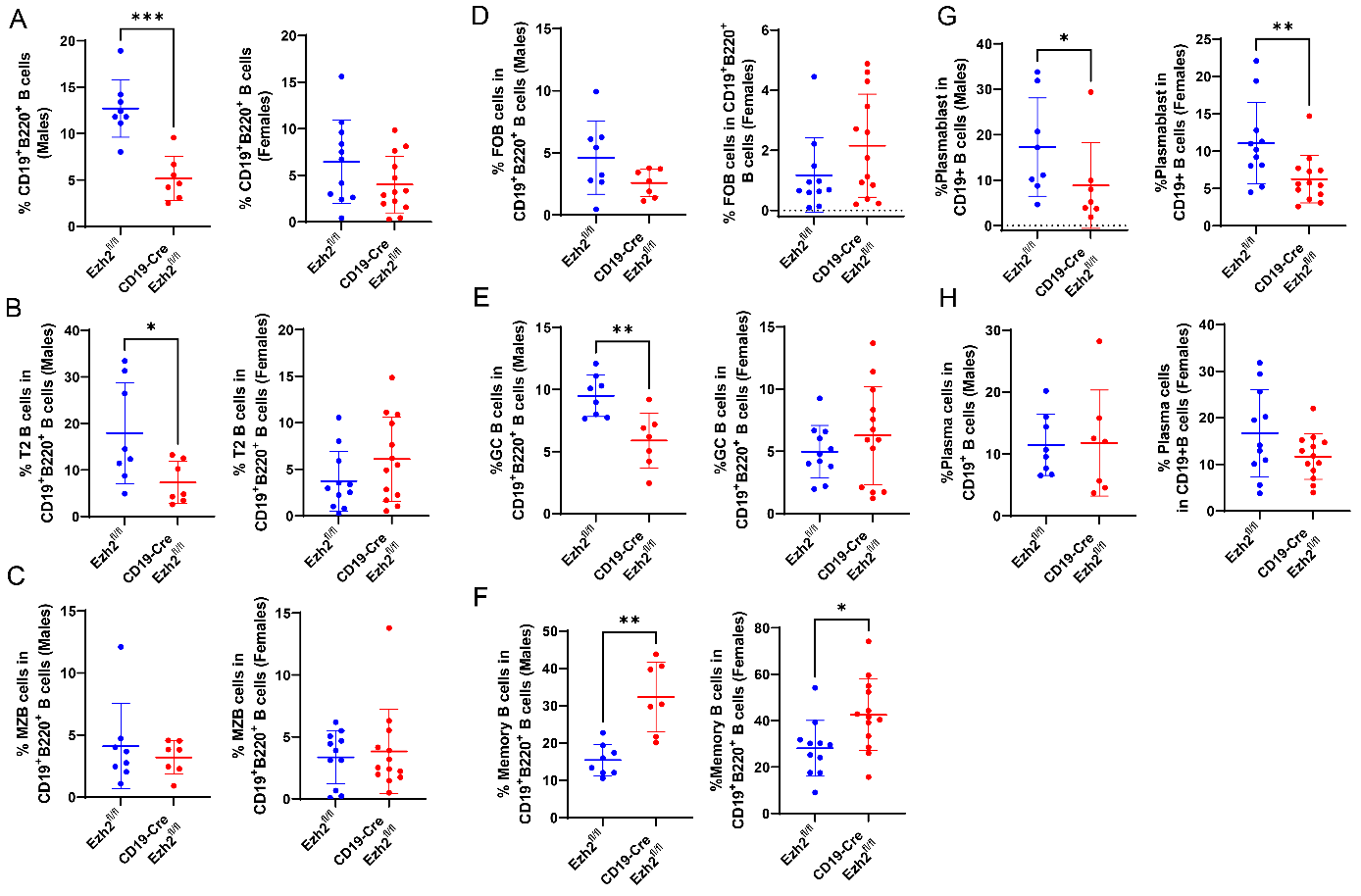
**Supplementary Figure 7. Sex-specific B cell development in the spleen of *Ezh2^fl/fl^* and CD19-Cre *Ezh2^fl/fl^* mice***.* Frequency of **(A)** CD19^+^B220^+^ B cells, and proportion of **(B)** transitional B cells (T2), **(C)** marginal zone B cells (MZB), **(D)** follicular B cells (FOB), **(E)** geminal center B cells (GC B), **(F)** memory B cells **(G)** plasmablast and **(H)** plasma cells in male mice (n=8 and n=7, respectively) and female mice (n=11 and n=13, respectively)in CD19^+^B220^+^ B cells in *Ezh2^fl/fl^* and CD19-Cre *Ezh2^fl/fl^* mice. * *p*<0.05, ** *p*<0.01, ****p*<0.001, two-tailed Mann-Whitney test.

**Supplementary Tables** (See Excel files)

**Supplementary Table 1.** Differentially expressed genes between clusters in single-cell RNA sequencing data of spleen B cells from *Ezh2^fl/fl^* and CD19-Cre *Ezh2^fl/fl^* mice.

**Supplementary Table 2.** List of differentially expressed genes in single-cell RNA sequencing of spleen B cells between *Ezh2^fl/fl^* and CD19-Cre *Ezh2^fl/fl^*.

**Supplementary Table 3.** Gene ontology (GO) analysis of biological process among genes differentially expressed between *Ezh2^fl/fl^* and CD19-Cre *Ezh2^fl/fl^* mice in both marginal zone B and germinal center B cell.
